# Co-administration of *Staphylococcus aures* Fibronectin Binding Protein A N1-3 region and Lipoteichoic acid antigen mixture enhance antibody response in BALB/c mice

**DOI:** 10.64898/2026.09.07.745799

**Authors:** Harish Babu Kolla, Rohini Krishna Kota, Sai Rupini Vemuri, Sai Archana Kurucherlapati, Prakash Narayana Reddy

## Abstract

*Staphylococcus aureus* (*S. aureus*) is a major human pathogen that causes a wide range of superficial skin infections to the life-threatening systemic diseases. Emergence and increasing prevalence of anti-biotic resistance *S. aureus* strains have accelerated the need for effective vaccines. As *S. aureus* pathogenesis involve its surface antigens to facilitate the colonization and invasion into host, targeting these surface antigens represent a promising strategy for *S. aureus* vaccine development. In this study, we evaluated the efficacy of two surface antigens of *S. aureus* -fibronectin-binding protein A (FnBPA) and lipoteichoic acid (LTA) as vaccine candidates. We expressed the N-terminal region of FnBPA in *Escherichia coli* and mixed with LTA to generate r-FnBPA N_1-3_+LTA antigen mixture, emulsified with Freund’s adjuvant as r-FnBPA N_1-3_+LTA antigen mixture or r-FR alone and immunized to BALB/c mice. Mice immunized with r-FnBPA N_1-3_+LTA mixture displayed a strong antibody response, strong bacterial growth inhibition *in vitro* and increased survival following lethal *S. aureus* challenge compared with mice immunized with r-FnBPA N_1-3_ alone. These findings suggest that incorporation of the *S. aureus* surface antigens like FnBPA with LTA provides improved protective efficacy and highlights the potential of a multi-antigen vaccine strategy for preventing *S. aureus* infections.

## Introduction

*Staphylococcus aureus* (*S. aureus*) remains as one of the most dangerous bacteria that infect humans and is responsible for various range of infectious diseases ranging from skin infections to life threatening endocarditis, sepsis, gastroenteritis, osteomyelitis and pneumonia [1–6]. Treatment has become very challenging due to the rapid evolutionary nature of the pathogen [7–8]. The emergence and wide spread of anti-biotic resistant strains like methicillin resistant *S. aureus* have exacerbated the global burden, posing severe challenges to the healthcare systems [9]. It is expected that the bacteria can evolve into more dangerous anti-biotic resistant strains in the coming years, making the treatment much more difficult.

Vaccination and immune therapy are effective ways to control and provide protection against the severity of disease. Despite decades of research on vaccine development, an effective *S. aureus* vaccine candidate has not yet been completely translated [10]. Continuous failure of the developed vaccines in the clinical trials highlight the complexity of *S. aureus* pathogenesis [11]. Therefore, the new immunoprophylactic strategies require inclusion of antigens that could elicit robust and durable immune responses. One of the defining characteristics of *S. aureus* pathogenesis is its ability to invade the host tissues through its surface associated adhesins which mediate the interactions with extracellular matrix proteins to facilitate immune evasion and promote persistence in diverse host niches.

One such structural component of *S. aureus* is Fibronectin binding protein A (FnBPA), which promotes its adhesion and invasion into host by mediating its interaction with host integrins via fibronectin [12–13]. Because of its role in the invasion and strong immunogenicity, FnBPA has been studied for its potential in vaccine development [14]. Parallelly, the lipoteichoic acid (LTA), a ubiquitous cell wall component in all the gram-positive bacteria [15], has been studied for its dual role in both bacterial pathogenesis and immune response. The LTA maintains the integrity of cell walls, formation of biofilms, and adhesion. It is recognized by the Toll-like receptor 2 (TLR2) [16–17]. Findings from the previous studies has shown that the anti-LTA antibodies mediate opsonophagocytic killing and thereby confer partial protection in the animal models. Nevertheless, LTA has not been investigated in combination of the protein antigens for *S. aureus* vaccine development, despite the potential for synergistic immune responses targeting both protein and non-protein bacterial components.

The main challenge in *S. aureus* vaccine development is to trigger immune response simultaneously interfering with adhesion, invasion, and survival in the host. Single-antigen strategies may not sufficiently neutralize the diverse virulence mechanisms employed by the pathogen, which suggests that rational combinations of antigens with complementary roles may provide superior protective efficacy. In this context, FnBPA and LTA represent a compelling pair: FnBPA as a key mediator of adhesion and invasion, and LTA as a structural surface component with immunostimulatory potential. By targeting both, we hypothesize that it may be possible to elicit antibody responses that block bacterial entry and enhance clearance.

To prove our hypothesis, we generated an immunogen mixture called r-FnBPA N_1-3_+LTA expressing N1-3 regions of FnBPA and then mixed with the LTA to give a protein-carbohydrate antigen conjugate. We evaluated the protective efficacy of the r-FnBPA N_1-3_+LTA mixture in comparison with the r-FnBPA N_1-3_ alone in the BALB/c mouse model. We identified that the administration of r-FnBPA N_1-3_+LTA immunogen mixture has shown robust antibody response, stronger inhibition of bacterial growth *in vitro* and increased the survival in mice challenged with the lethal dose of *S. aureus* bacteremia as compared to the r-FnBPA N_1-3_ alone. Therefore, our findings highlight that the combination of protein and non-protein surface antigens may provide a better immune response and overcome the limitations of single antigen-based vaccines and support the rationale for advancing such strategies in the development of broadly effective *S. aureus* vaccines.

## Materials and Methods

### Chemicals, reagents and media

The dehydrated media, antibiotics and other supplements were procured from HiMedia, India. Mouse anti-6X-Histidine monoclonal antibodies and HRP-conjugated secondary antibodies and Lipoteichoic acid (L3140) purified (≥90 % pure) from *Streptococcus pyogenes* were purchased from Sigma-Aldrich, India. Biotin-3-sulfo-N-hydroxysuccinimide ester sodium salt (B5161) is from Sigma. The pET22b plasmid, cloning and expression hosts *Escherichia coli* NovaBlue and BL21 (DE3) strains were from Novagen.

### Animals

The BALB/c mice were caged in polypropylene cages bedded with rice hulls and acclimatized to 12 hours day-night cycle in the in-house animal house facility. The cages were cleaned and changed every 2-3 days. used in this study were maintained in 12-h light and dark cycles alternately. Mice were provided with food pellets and mineral water from throughout the study period.

### Cloning and expression

Initially, the nucleotide portions of N1-3 region in FnBPA were PCR amplified and cloned into pET22b vector between the BamHI and HindIII restriction sites as like our previous studies [18–21]. The recombinant plasmid was transformed into *E. coli* NovaBlue strain and the transformants were screened through colony PCR using the T7 primers. The confirmed transformants were further propagated followed by plasmid isolation and subsequent transformation into the expression host *E. coli* BL21(DE3) pLysS (Novagen, USA). Around 2 mL of Overnight culture of *E. coli* BL21(DE3) pLysS carrying pET22b-r-FnBPA N_1-3_ recombinant plasmid was added to around 200 mL of Luria–Bertani broth supplemented with 100 μg/mL of ampicillin and 35 μg/mL of chloramphenicol for the recombinant protein expression. The overnight culture was induced with the 1 mM of isopropyl β-D-galactopyranoside (IPTG) and incubated for another 5 h. After this, the cells were harvested by centrifuging the culture at 9000 g followed by lysis. The cells were lysed, and the lysate was centrifuged at 25000g at 25°C to separate the aqueous protein phase from the LPS-rich detergent layer to prevent the LPS contamination in the protein. The recombinant protein was further purified under native conditions from periplasmic extracts using imidazole and Ni–NTA resin by immobilized metal affinity chromatography (IMAC) (Qiagen, Germany) where the non-recombinant proteins, cell debris and contaminants like LPS were washed off during the washing processes. The concentrated recombinant protein was buffer exchanged in 1 × PBS and the concentration was determined by Lowry’s colorimetric assay using BSA as standard as we followed in our previous studies [18–21].

### Western Blot

Around 20 ug of the purified protein was separated on 12 % SDS-PAGE gels in tris–glycine buffer. The separated protein was transferred on the nitrocellulose membrane by wet-transfer method using Bio-Rad transfer apparatus. The blots were blocked with the 5 % skim milk followed by subsequent washing and then incubated with the 1: 1000 diluted serum obtained from mouse immunized with the r-FnBPA N_1-3_ diluted in the 1XPBST buffer. The blots were washed properly with the 1XPBST buffer the next day and probed with the 1: 2000 diluted secondary anti-mouse antibody conjugated with HRP for 45 mins. The blots were developed by adding a pinch of diaminobenzidine tetrahydrochloride (DAB) (Sigma-Aldrich, India) in 0.03% H_2_O_2_ in PBS.

### Antigen Preparation and Immunization

The antigen was prepared in the complete or incomplete Freund’s adjuvant for the immunization in female BALB/c mice. The antigens selected for immunization were as follows: PBS control; purified recombinant r-FnBPA N_1-3_ protein and r-FnBPA N_1-3_ along with the LTA (r-FnBPA N_1-3_+LTA). For the immunization, around five-week-old female BALB/c mice (n=6 each group) were categorically divided into three groups receiving PBS (hereafter referred as Sham); r-FnBPA N_1-3_ and r-FnBPA N_1-3_+LTA. Around 50 ug of the purified r-FnBPA N_1-3_ or r-FnBPA N_1-3_+LTA complexed with 100 ug of LTA per 200 uL per mouse for 1 hr at room temperature was emulsified with the complete Freund’s adjuvant and administered subcutaneously to the mice. While the Sham group received PBS with complete Freund’s adjuvant. Following this, two booster doses were administered with the same antigen emulsified with Freund’s incomplete adjuvant for the second and third doses at 14^th^ and 28^th^ days respectively. Blood was collected for serum collection every week through tail vein puncture. The antibody levels were determined by indirect ELISA using recombinant r-FR as antigen.

### Bacterial Growth Inhibition Under the Tested Conditions

To determine the anti-bacterial activity of the serum obtained from r-FnBPA N_1-3_ and r-FnBPA N_1-3_+LTA immunized mice against the *S. aureus*, we performed the *in vitro* anti-microbial assay in a 96 well plate as triplicates. Approximately 200 uL of the overnight culture of *S. aureus* NCIM 2127 was added to each well and around 50 ug of the diluted serum (sham, r-FnBPA N_1-3_ and r-FnBPA N_1-3_+LTA) was added to the corresponding wells and incubated at 37°C for 12 hours. Absorbance at 600 nm was recorded 12 hours after seeding. The extent of bacterial growth inhibition by the serum was then determined based on the final OD values.

### Animal Challenge Studies

The protective efficacy of passive administration of immune sera from all the three groups was evaluated in the BALB/c mice with *S. aureus* challenge. The mice were divided into three groups, each group receiving around 250 uL of ten-fold diluted anti-Sham; anti-r-FnBPA N_1-3_ and anti-r-FnBPA N_1-3_+LTA serum collected from the actively immunized animals previously. The serum was administered intravenously and left unchallenged for 24 hours. After 24 hours, the animals are challenged with the lethal dose of 2.5 × 10^7^ CFU and monitored for 10 days. The bacterial load was further confirmed through plating and then proceeded for the challenge studies. For the challenge of lethal dose, tubes were labelled for each mice receiving 2.5 × 10^7^ CFU and the fresh overnight culture of bacteria was injected to each mouse through intravenous route [22]. Deaths in each group over these 10 days period were recorded for calculating the survival percentage.

### Epitope prediction

The N-terminal end of FnBPA protein was used to predict both the B and T cell epitopes. For the linear B cell epitope prediction, the amino acid sequence was submitted to the BepiPred 2.O (https://services.healthtech.dtu.dk/services/BepiPred-2.0/) [23] and B cell epitopes were predicted using the default settings. The peptides with more than the threshold score of 0.5 were considered epitopes. Whereas for the prediction of conformational B cell epitopes, the protein structure was predicted through protein modeling, and the refined protein structure was submitted to the Discotope tool in the IEBD server (http://tools.iedb.org/discotope/) [24]. The CD8+ T cell epitopes were predicted using the IEDB server corresponding to the BALB/c MHC I alleles K^d^, D^d^ and L^d^ respectively (https://www.iedb.org/). The peptides with the IC50 values less than 500 nM were selected as CD8+ T cell epitopes [25]. Similarly, The CD4+ T cell epitopes were predicted with the cut-off IC50 values of 1000 nM for the MHC II alleles H-2-IA^d^ and H-2-IE^d^ [23].

### Statistical analysis

The data presented in this manuscript are represented as mean±SD. All statistical analyses were performed using one-way ANOVA in GraphPad Prism version 10.3.1. P values were considered statistically significant as follows: *P < 0.05, **P < 0.01, ***P < 0.001, ****P < 0.0001.

## Results

### Epitope prediction

*In silico* analysis of the N1-3 segments of FnBPA has shown that the selected antigen from the N terminal domain of this protein is immunodominant and rich in the B cell epitopes and able to trigger a strong humoral immune response. We found a total of 12 linear B cell epitopes (highlighted with different colours in the amino acid sequence) in the N1-3 segments of FnBPA (**Figure 1A**) (**Table 1**) indicating its ability to generate and reactive to a wide range of polyclonal antibodies. Structural analysis has shown the presence of conformational B cell epitopes in the FnBPA N1-3. Almost 40 % of the residues fall under the positive prediction of conformational B cell epitopes highlighting their immunodominant nature to elicit a string B cell mediated immunity (**Figure 1B**). The T cell epitope data has also shown that the N1-3 segments of FnBPA are rich in both the CD8+ and CD4+ T cell epitopes (**Figure 2** (**Table 1**). We predicted a total of 3 CD8+ epitopes and 10 CD4+ T cell epitopes highlighting that the antigen can elicit a strong cell mediated immune response as well (**Figure 2A and 2B**) (**Table 1**). One interesting finding we observed was that the presence of adjacent or overlapping CD4+ T cell epitopes. There is a total of 3 regions in the FnBPA N1-3 where the CD4+ T cell epitopes are overlapping-TQVEVAQPRTASESKPRVTRSADVAEAKEA (TQVEVAQPRTASESK and PRVTRSADVAEAKEA); KANNRFSHVAFIKPNNGKTTSVTVTGTLMK (KANNRFSHVAFIKPN, FSHVAFIKPNNGKTT and NGKTTSVTVTGTLMK); and EDIAKSVYANTTDTSKFKEVTSNMSGNLNL (EDIAKSVYANTTDTS and KFKEVTSNMSGNLNL) (**Figure 2B**).

**Figure 1.**
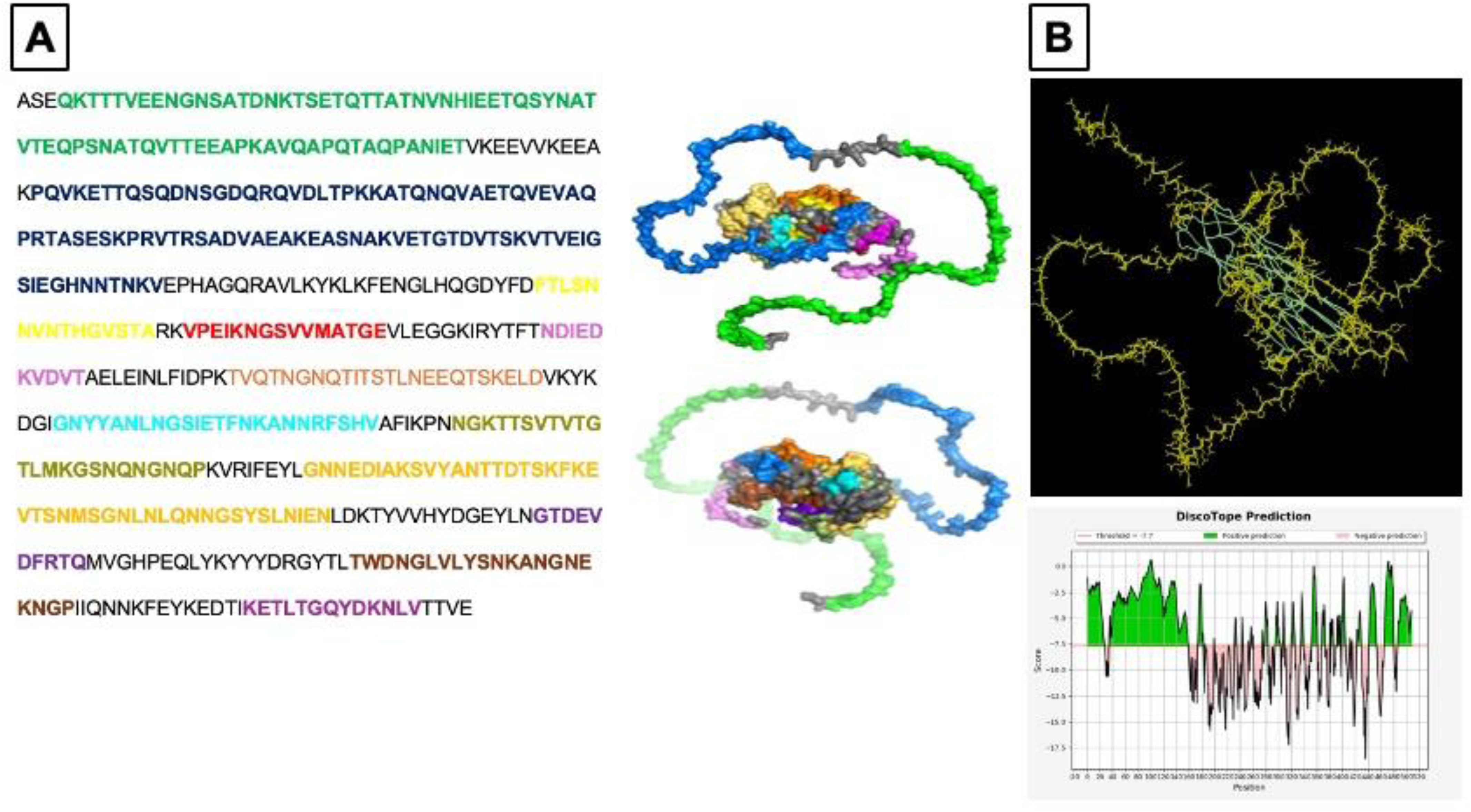
**A.** linear B cell epitopes and **B.** conformational B cell epitopes in the FnBPA N1-3 antigen.

**Figure 2.**
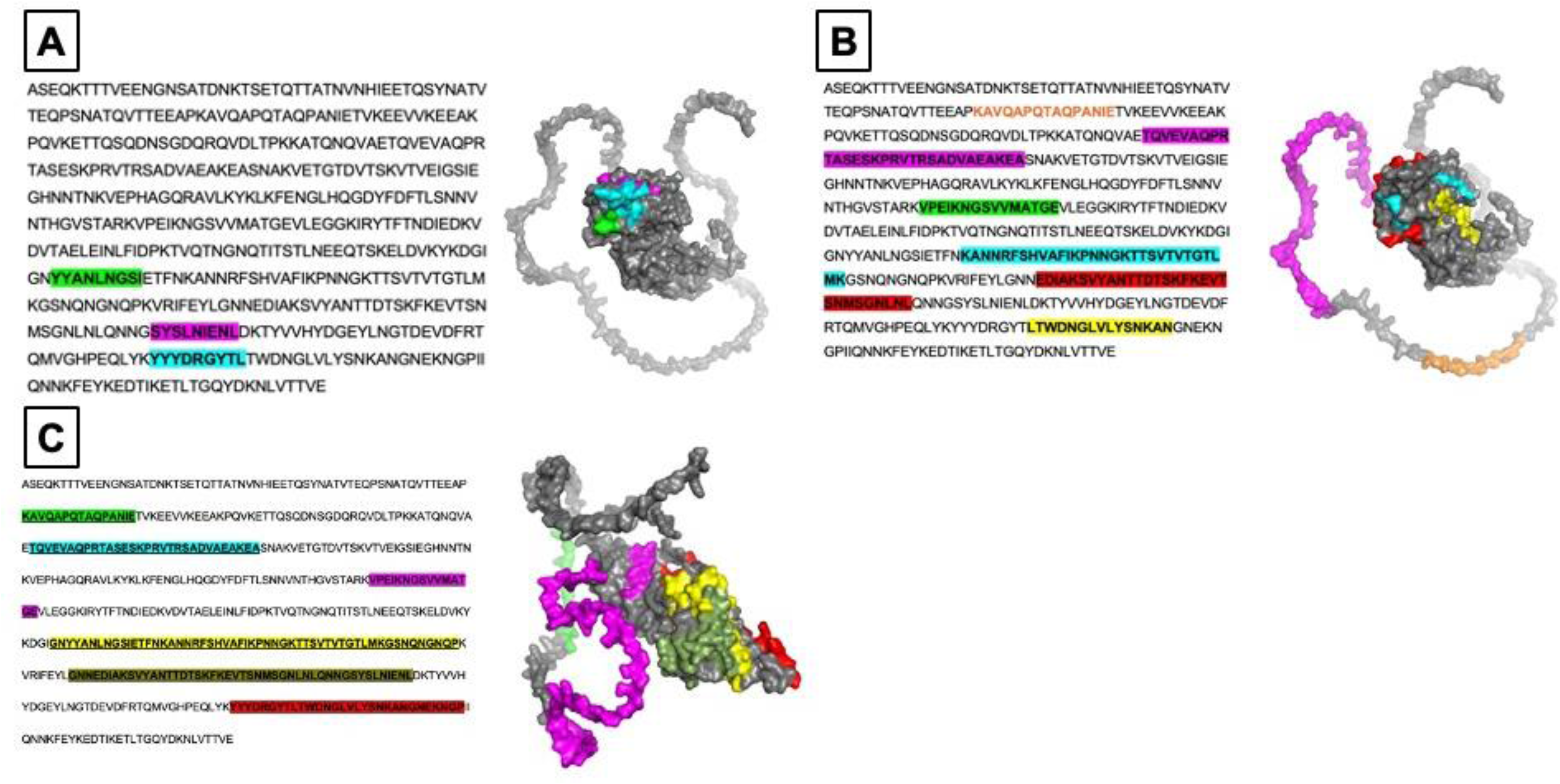
**A.** CD8+ and **B.** CD4+ T cell epitopes in the FnBPA N1-3 antigen. **C.** Immunodominant regions showing the overlapped B and T cell epitopes in the antigen.

**Table 1.**
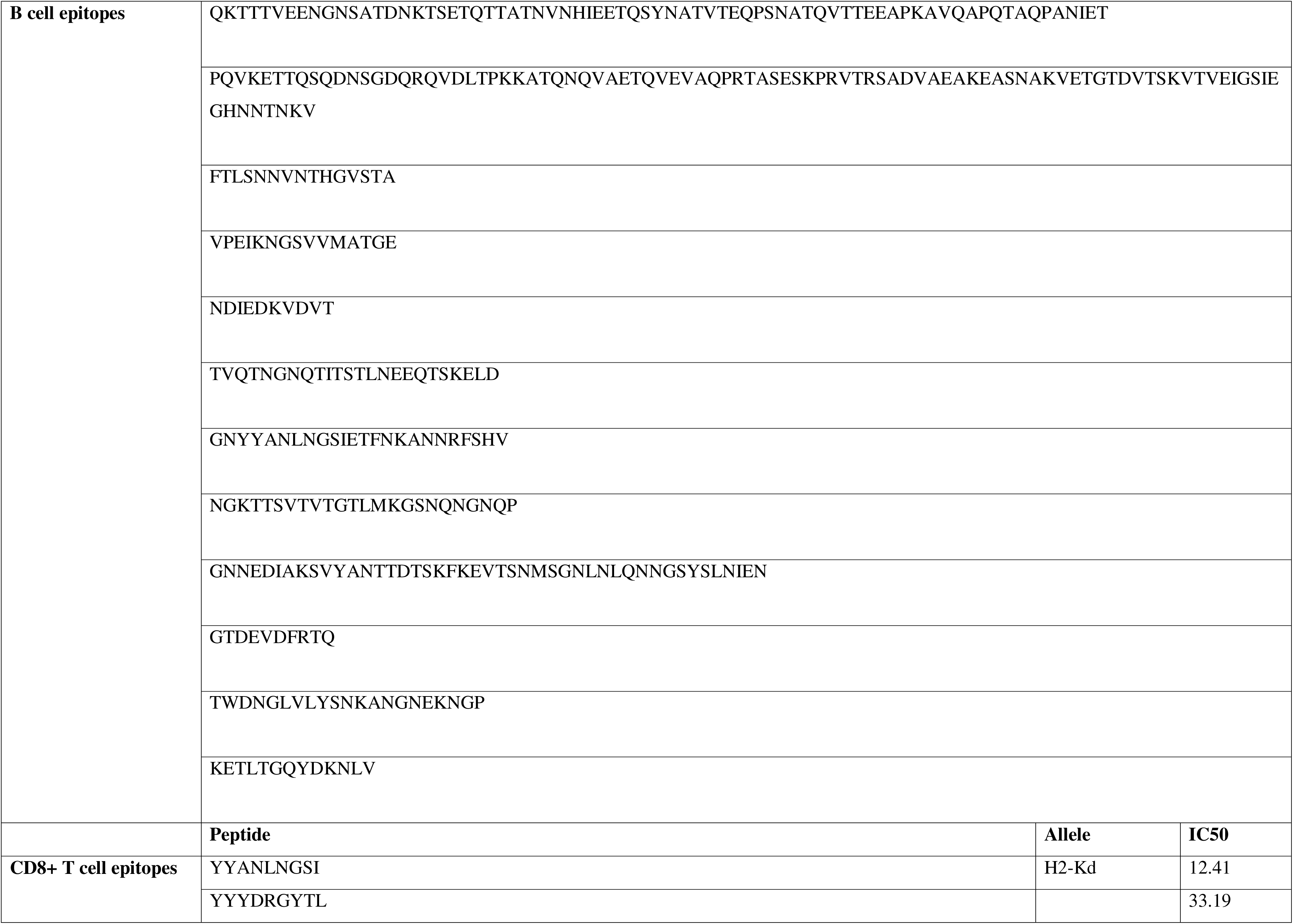
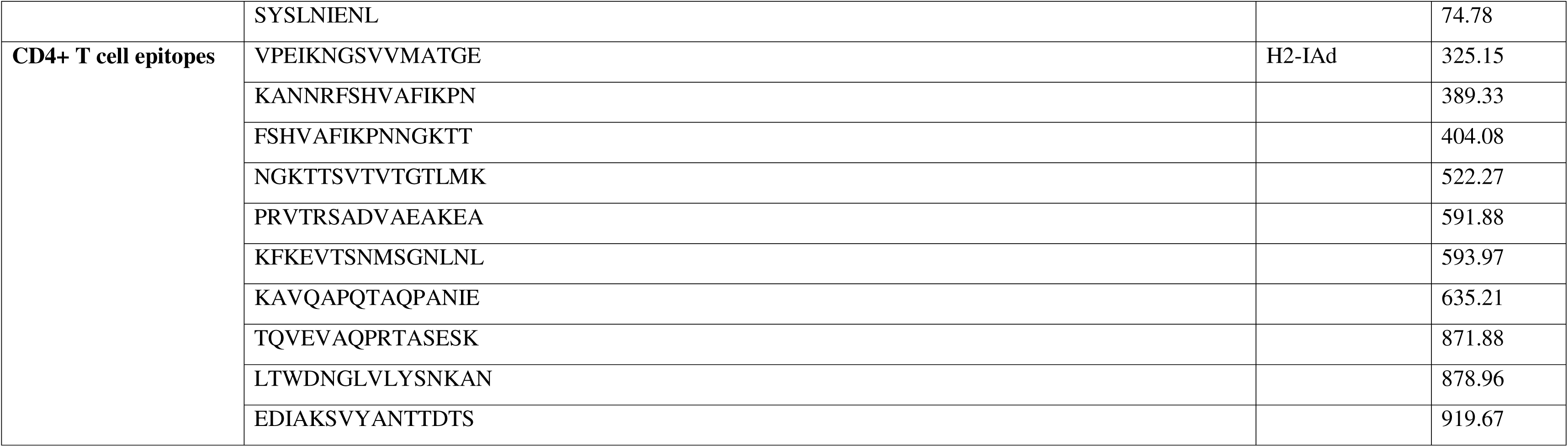
List of B and T cell epitopes predicted in the FnBPA N1-3 antigen.

### Immunodominant peptide regions

We also observed that the B and T cell epitopes (both CD8+ and CD4+) also overlapped together forming 6 immunodominant oligopeptide regions in the FnBPA N1-3 antigen (**Figure 2C**). Firstly, the peptide region KAVQAPQTAQPANIE is a CD4+ T cell epitope which is an integral part at the C-terminal end a B cell epitope QKTTTVEENGNSATDNKTSETQTTATNVNHIEETQSYNATVTEQPSNATQVTTEEAPKAVQAPQTAQPANIET. Similarly, the peptide region TQVEVAQPRTASESKPRVTRSADVAEAKEA comprises of an overlapping CD4+ T cell epitopes QVEVAQPRTASESK and PRVTRSADVAEAKEA as discussed in the above section, is also part of a B cell epitope PQVKETTQSQDNSGDQRQVDLTPKKATQNQVAETQVEVAQPRTASESKPRVTRSADVAEAKEASNAKVETGTDVTSKVTVEIGSIEGHNNTNKV. The CD4+ T cell epitope VPEIKNGSVVMATGE was also predicted to be a linear B cell epitope which possesses the ability to induce both the cell mediated and humoral immune response.

The peptide GNYYANLNGSIETFNKANNRFSHVAFIKPNNGKTTSVTVTGTLMKGSNQNGNQP comprises of 2 linear B cell epitopes GNYYANLNGSIETFNKANNRFSHV, NGKTTSVTVTGTLMKGSNQNGNQP, a CD8+ T cell epitope YYANLNGSI and 3 CD4+ T cell epitopes KANNRFSHVAFIKPN; FSHVAFIKPNNGKTT; and NGKTTSVTVTGTLMK. Additionally, one B cell epitope GNNEDIAKSVYANTTDTSKFKEVTSNMSGNLNLQNNGSYSLNIEN,a CD8+ T cell epitope SYSLNIENL And two CD4+ T cell epitopes EDIAKSVYANTTDTS and KFKEVTSNMSGNLNL constitute a long oligopeptide region GNNEDIAKSVYANTTDTSKFKEVTSNMSGNLNLQNNGSYSLNIENL. Finally, the peptide region YYYDRGYTLTWDNGLVLYSNKANGNEKNGP covers a B cell epitope TWDNGLVLYSNKANGNEKNGP, a CD8+ T cell epitope YYYDRGYTL, and a CD4+ T cell epitope LTWDNGLVLYSNKAN.

### Cloning and expression of recombinant N1-N3

The N1-3 segments of the N terminal end of FnBPA protein were PCR amplified with the BamHI restriction overhang at 5’ end and HindIII restriction site at the 3’ end resulting in a 1500 bp long fusion gene r-FnBPA N_1-3_ (**Figure 3A-B**) and hetelogously expressed in the *E. coli* BL21DE3 strain with IPTG induction. Recombinant expression of the r-FnBPA N_1-3_ protein was purified, separated on 12 % gel and detected at the desired molecular weight of ∼50KDa with serum from both r-FnBPA N_1-3_ immunized mice (**Figure 4A-B**).

**Figure 3.**
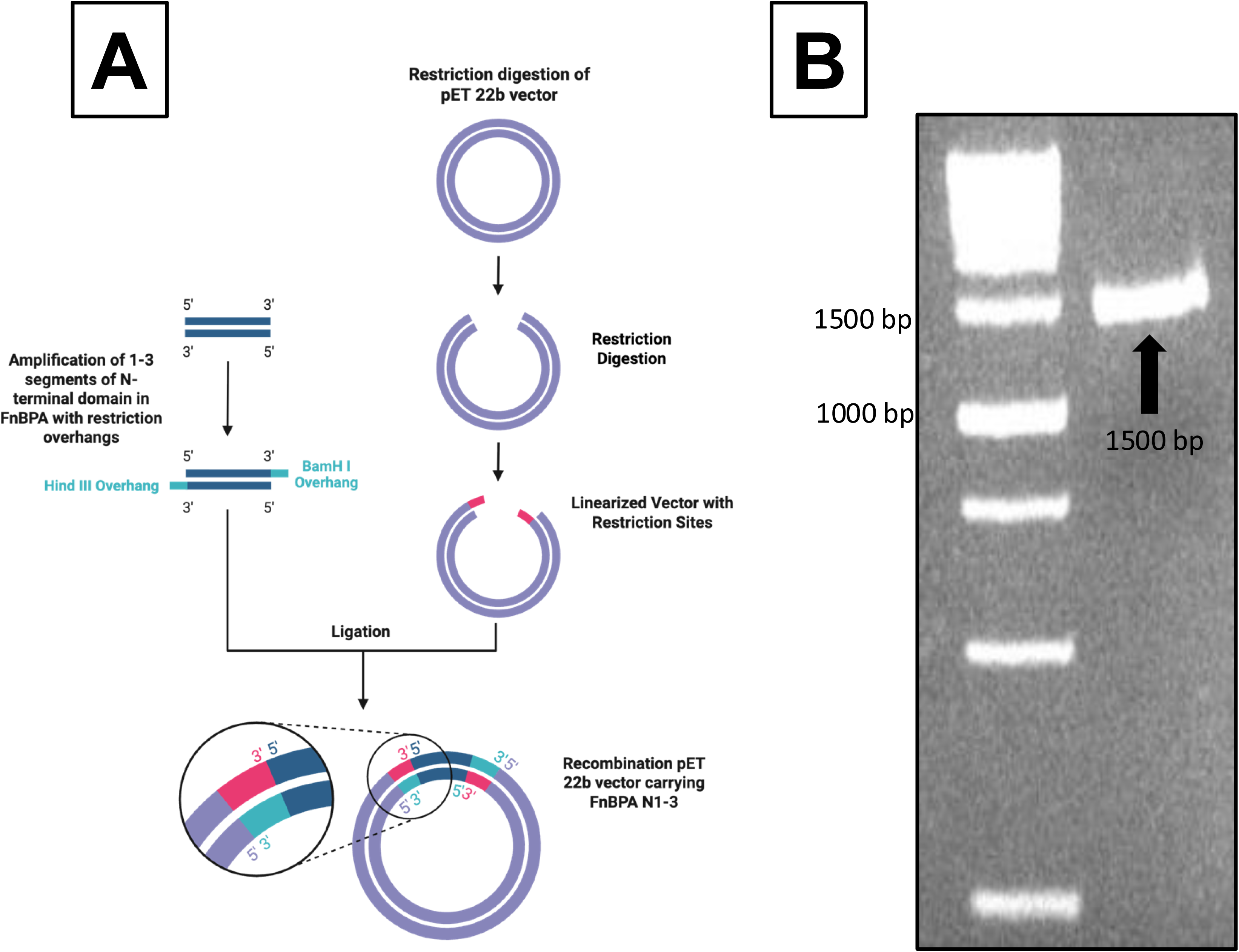
Cloning and expression of FnBPA N1-3 gene region. **A.** Schematic representation of the cloning strategy. **B.** Agarose gel image of FnBPA N1-3 amplicon; **C.** SDS-PAGE image of purified r-FR.

**Figure 4.**
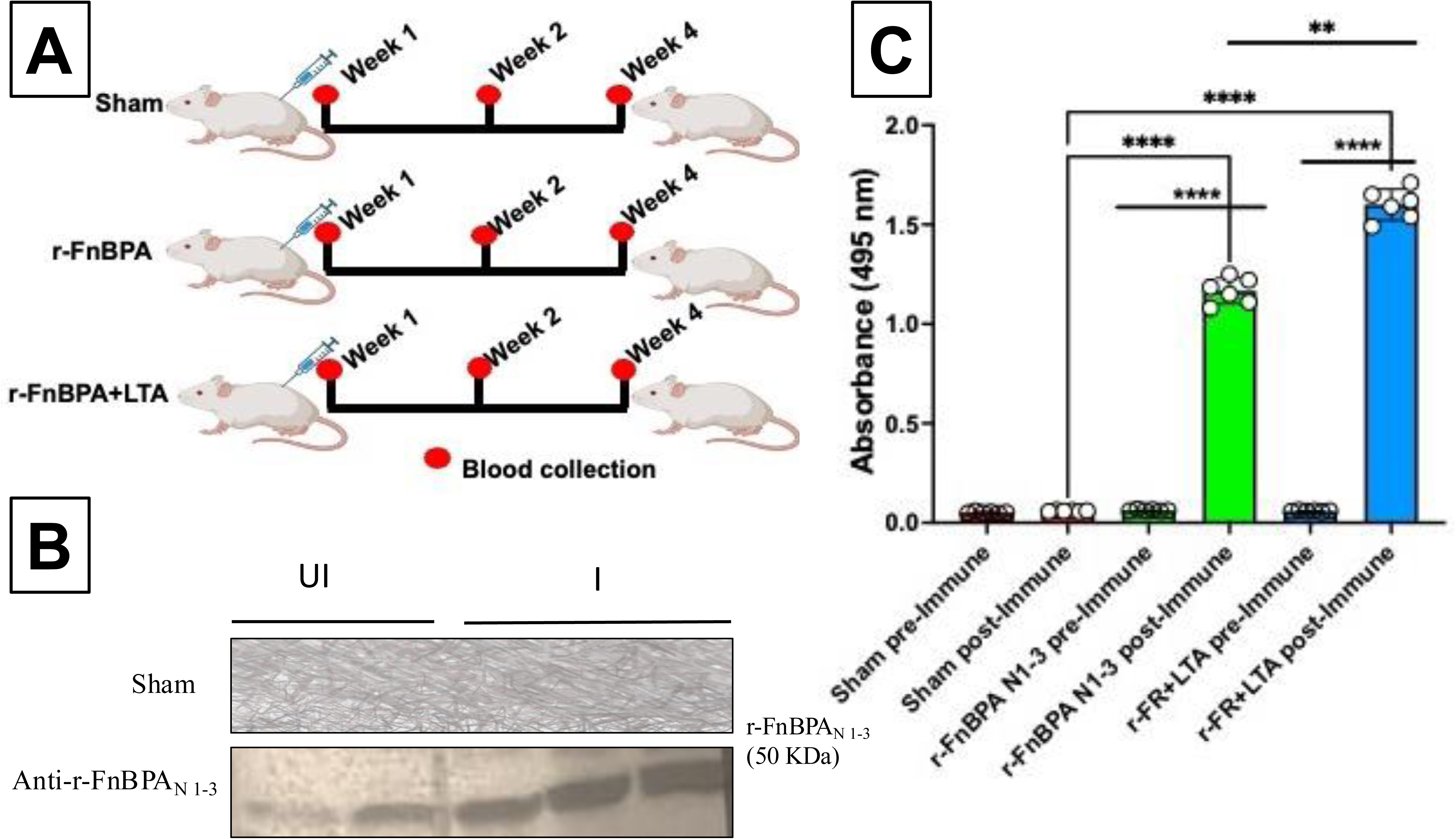
Immunization of r-FnBPA_N 1-3_ to the BALB/c mice. A. Schematic representation of the animal study. B. Western blot images showing the seroreactivity of anti-r-FnBPA_N 1-3_ serum against the recombinant protein, C. Bar graphs showing the antibody levels in the three groups (Sham vs r-FnBPA_N 1-3_ vs r-FnBPA_N 1-3_+LTA). All the data represented in mean ± SD (n=6). P values were considered statistically significant as follows: *P < 0.05, **P < 0.01, ***P < 0.001, ****P < 0.0001.

### Antibody responses

We determined the antibody levels in all the three immunized groups. Blood was collected from all the three groups pre-immunization to compare the antibody levels with post-immunization. We observed that there is no appreciable difference in the total antibody levels with Sham immunization before and after immunization, while the antibody levels increased with r-FnBPA N_1-3_ and r-FnBPA N_1-3_ +LTA groups post-immunization (**Figure 4C**). Overall, the total antibody levels in r-FnBPA N_1-3_ +LTA group post-immunization are greater than r-FR alone, making a significant difference (**Figure 4C**).

### Microbial Growth Inhibition Under Testing Conditions

We later assessed the anti-bacterial activity of serum under controlled conditions *in vitro* obtained from actively immunized animals to see if the post-immune serum can inhibit bacterial growth *in vitro*. The bacterial growth inhibition was seen with the post-immune serum in both the groups whereas the inhibition is more prominent with the anti-r-FnBPA N_1-3_ +LTA sera even at the same concentration (**Figure 5A**). After 12 hours of culturing, the anti-bacterial activity was significant in the anti-r-FnBPA N_1-3_ and anti-r-FnBPA N_1-3_ +LTA groups based on the final OD values (**Figure 5A**).

**Figure 5.**
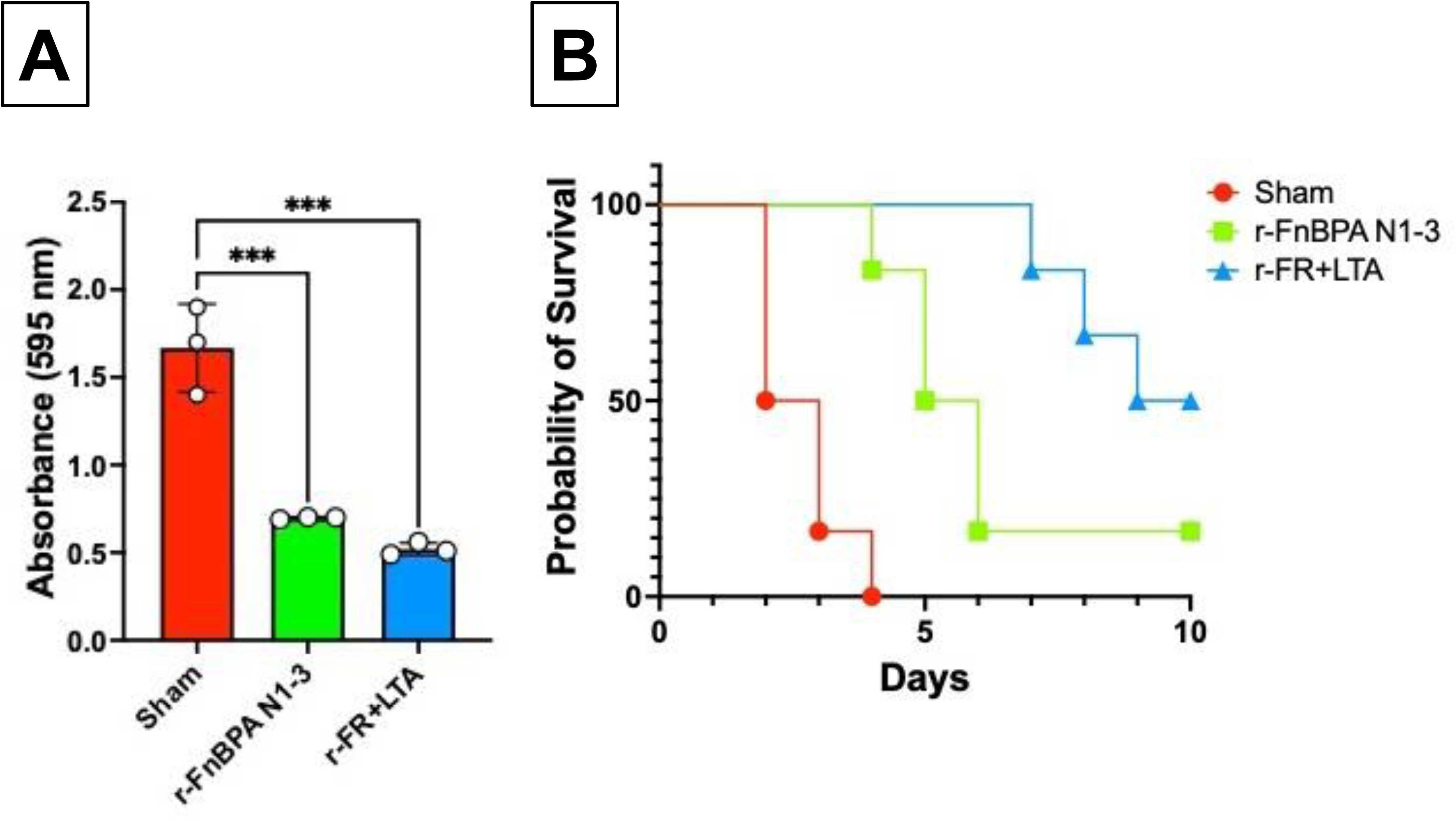
r-FnBPA_N1-3_+LTA exhibited potent bacterial growth inhibition and passive protective *in vivo*. A. Bacterial growth inhibition after 12 hours. B. Kalpan-meier curve shows the passive protective effect of antibodies from r-FnBPA N_1-3_ and r-FnBPA N_1-3_+LTA immunizations. P values were considered statistically significant as follows: ***P < 0.001.

### Protective Capacity of Immune Sera

The protective efficacy of r-FR and r-FR+LTA anti-sera was assessed through animal challenge studies for a period of 10 days. The animals were closely monitored and the total number of deaths in each group was recorded. In the case of control group that were challenged with the lethal dose of *S. aureus* and had received the serum from Sham immunized group, all the 6 mice were dead within 4 days (**Figure 5B**). While the mice received anti-FR serum has shown a survival rate of ∼16.66 % after 10 days where one mouse survived the challenge, and the others were dead within 6 days. Importantly, in the animals received anti-r-FR+LTA serum, the deaths were reported lately on day 7 (n=1), 8 (n=1) and 9 (n=1) as compared to the other two groups (**Figure 5B**). Therefore, the overall survival rate of this group is greater than the other two with almost 50 % (**Figure 5B**). Overall, the animal challenge studies have shown that the anti-r-FR+LTA sera has shown a better protective capacity in the animals than r-FR alone.

## Discussion

In this current study, we developed a novel conjugate vaccine candidate r-FnBPA N_1-3_+LTA against the *S. aureus* by expressing the N1-3 regions of FnBPA and then mixed with the LTA, a bacterial cell wall polymer. Our findings highlight that the r-FnBPA N_1-3_+LTA elicits a strong humoral immune response and protection against the *S. aureus* strain in mouse model as compared to the single antigen immunization with the r-FnBPA N_1-3_ comprising of FnBPA N1-3 region. In brief, the vaccine candidate induced strong antibody responses which were determined by comparing the pre and post immune sera in all the three groups (Sham, r-FnBPA N_1-3_and r-FnBPA N_1-3_+LTA). The antibody rich post-immune sera demonstrated a strong anti-bacterial against the *S. aureus* cultures underscoring the promise of multi-antigen vaccine strategies in combating *S. aureus* infections.

These results are significant given the disappointing track record of single antigens in the prior *S. aureus* vaccine candidates [10, 26]. In this context, the r-FnBPA N_1-3_+LTA strategy—combining conserved adhesin domains (FnBPA N1–3) with a highly immunogenic cell wall component LTA addresses the need for broader, multi-epitope coverage. The simultaneous presentation of protein and non-protein antigens to the host immune system may better mimic natural bacterial surfaces, leading to functionally protective immune responses rather than non-neutralizing antibodies. Most importantly, the polyclonal antibodies generated by r-FnBPA N_1-3_+LTA have not only exhibited broad anti-microbial activity but also conferred passive protection against the lethal dose of *S. aureus* as compared to r-FnBPA N_1-3_. This ability of the bivalent fusion conjugate r-FnBPA N_1-3_ +LTA differentiates it from the single antigen candidates alone.

Besides the enhanced antibody response with r-FnBPA N_1-3_+LTA, we acknowledge that our experimental design does not exclude the possibility of non-specific adjuvant effects of LTA like activation of innate immune pathways, cytokine production and antigen-presentation. Hence, we expect some extent of adjuvant-driven enhancement in immune response independent of scaffold conjugation. However, our interpretation underscores that the conjugated LTA presentation may enhance the efficiency of antibody production, rather than claiming that the scaffold alone is solely responsible for the improved outcome.

We acknowledge several limitations in our study. Firstly, this report comprises of the epitope prediction and these findings do not incorporate the experimental validation of peptides and biomolecular interactions linking these computational findings to functional immune outcomes such as cytokine secretion, T-cell activation, opsonophagocytic killing, or complement-mediated activity. On the other hand, lack of comprehensive cytokine profiling, T cell mediated immunity studies other mechanistic studies represents a significant limitation. While these functional studies are beyond the scope of our study as our primary interest lies on the antibody response and microbial inhibition with the multiple surface antigen presentation approach under the controlled settings. However, our computational predictions and the preliminary findings provide a biologically plausible framework for immune engagement and serve as a foundation for subsequent experimental validation. Accordingly, our findings should be interpreted as hypothesis-generating rather than definitive evidence of protective immunity.

Moreover, we used BALB/c model because of several reasons like the availability and reproducibility of reagents to study immune responses. But this model does not completely replicate the native *S. aures* conditions in humans for the translational applications. Further studies in various human cell lines and organoid models are of high importance prior to its clinical trials [27–30]. Further validation of the vaccine candidate in conferring protection in multiple preclinical models—including bloodstream, pneumonia, and device-associated infections are also essential. Additionally, safety considerations are also important. Although the LTA is highly conserved and potential antigen, it triggers pro-inflammatory response by activating the TLR2 [31]. Excess inflammatory responses must be carefully examined, and the dose of LTA needs to be properly optimized for the translational studies.

## Conclusion

In conclusion, this study highlights the potential of r-FnBPA N_1-3_+LTA antigen mixture as a next-generation *S. aureus* vaccine candidate. By addressing the shortcomings of previous monovalent approaches and leveraging a multi-antigen, structurally stable design, r-FnBPA N_1-3_+LTA offers a rational and promising path toward an effective prophylactic strategy. Given the rising threat of MRSA and the global crisis of antibiotic resistance, advancing such candidates into further preclinical and translational studies is both timely and urgently needed. The mice immunized with r-FnBPA N_1-3_+LTA elicited higher antibody response, enhanced inhibition of microbial growth *in vitro*, and protective capacity of sera against the pathogen challenge in BALB/c mouse model as compared to the fusion protein r-FnBPA N_1-3_ alone. This study highlights the importance of surface antigens in *S. aureus* vaccine development.

## Acknowledgement

PNR thanks Department of Science and Technology for awarding the DST-INSPIRE Faculty award and RKK thank Council of Scientific and Industrial Research for providing senior research fellowship during the study period.

## Author contributions

HBK, RKK, and PNR performed the experiments. HBK, SRV, and SAK analyzed the data. HBK and PNR wrote the manuscript. All authors have approved the final version of the manuscript for publication.

## Data availability

All data generated or analyzed in this study are included in this article. The raw data will be provided upon request made to the correspondent author.

## Declaration

### Ethical approval

All the experiments carried out in the current study were according to the guidelines from Institutional Animal Ethics Committee, of Vignan’s Foundation for Science, Technology, and Research (Reg. No.: 2046/PO/ReBi/S/18/CPCSEA), Guntur, Andhra Pradesh, India.

### Consent to participate

Not applicable

### Consent to publication

Authors declare that there is no conflict of interest.

